# Comparison of urinary proteome in the first two days after mating in male rats

**DOI:** 10.1101/2023.11.18.567698

**Authors:** Haitong Wang, Chenyang Zhao, Youhe Gao

**Author notes:** Corresponding author: Gao Youhe (1964.06 -), male, professor, doctoral supervisor. His research interests include urine proteomics and urine biomarkers. **Fund Project:** Beijing Normal University (11100704). **About the author:** 1. Wang Haitong. (2000.4 -), female, master candidate. Her research interests include urine biomarkers. 2. Zhao Chenyang. (Second Zuo) (1998.03 -), female, master candidate. Her research interests include urine biomarkers.

## Abstract

**Objective:** To explore whether differences between male rats on the next day of mating and on the day of mating can be reflected by the urine proteome.

**Methods:** Urine samples were collected from male Sprague-Dawley rats on the day of mating and the next day of mating. Urine samples were analysed by the free-labelled quantitative proteomics technique of high-performance liquid chromatography tandem mass spectrometry (LC-MS/MS). Differential proteins of the urine proteome were analysed for protein function and biological pathways.

**Results:** 43 differential proteins were identified by comparing the urine proteome of rats on the next day of mating with that on the day of mating, and nearly two-thirds of the differential proteins were related to spermatogenesis.

**Conclusions:** The urine proteome has the potential to reflect spermatogenesis without interfering with it.

## 1 Introduction

Spermatogenesis is the process by which spermatogonial cells undergo proliferation and differentiation to form mature sperm, which involves chromosome fold reduction and cell deformation. There are three stages of mammalian spermatogenesis. First, the spermatogonium undergoes a series of mitosis to form a primary spermatocyte. Secondly, the primary spermatocytes produce haploid round sperm cells through meiosis. Finally, the round sperm cells are deformed into sperm with flagella. Paracrine, autocrine, and endocrine pathways all contribute to the regulation of this process, and the large number of structural elements and chemical factors involved in regulation make the network connecting various cellular activities during spermatogenesis unimaginably complex^1^. Urine is produced by the filtration of blood through the kidneys to eliminate metabolic waste, which is not controlled by the homeostatic regulation mechanism of the internal environment, and can be more sensitive to the various small changes produced by the body^2^. It has been shown that the urine metabolome can be used to differentiate normozoospermic infertile men from fertile individuals^3^. However, no studies have been conducted to monitor spermatogenesis by urine proteome. After a mating of male rats, mature sperm is consumed to stimulate spermatogenesis in the testis. In this study, by collecting urine samples from rats on the day of mating and the next day of mating, we compared the urinary proteome and tried to explore whether spermatogenesis could be reflected in the urinary proteome.

### 2.1 Experimental material

#### 2.1.1 Experimental animal

Five 10-week-old male Sprague-Dawley rats and five 10-week-old female Sprague-Dawley rats were purchased from Beijing Weitonglihua Experimental Animal Biotechnology Company Limited. All rats were raised in a standard environment (room temperature 22±1 °C, humidity 65%-70%). All the rats were fed in the new environment for three days before the experiment began. All experimental operations were reviewed and approved by the Ethics Committee of the College of Life Sciences, Beijing Normal University with the approval number CLS-AWEC-B-2022-003.

### 2.2 Experimental method

#### 2.2.1 Rats mating

Each of the five rats was cohabited with a female rat at 16:00. At 7:00 the next day, female Rats were checked for vaginal plugs, and those who detected vaginal plugs were considered to be mating between male and female.

#### 2.2.2 Urine sample collection

On the mating day, each of the five rats was placed into a single metabolic cage from 20:00 to 8:00 the next day and urine was collected through the metabolic cage. Urine was collected separately and temporarily stored in the -80□ refrigerator as urine samples on the mating day. After urine collection, male rats were fed separately. On the next day after mating, each of the five rats was placed into a single metabolic cage from 20:00 to 8:00 the next day and urine was collected through the metabolic cage. Urine was collected separately and temporarily stored in the -80□ refrigerator as urine samples on the next day after mating.

#### 2.2.3 Urine sample treatment

Urine protein extraction: Rat urine samples were removed from the refrigerator at -80 °C and thawed at 4°C. Rat urine samples were centrifuged at 4°C, 12000×g for 30 min and the supernatants were transferred to new Eppendorf (EP) tubes. Triple volumes of precooled absolute ethanol was added, and the samples were homogeneously mixed and precipitated overnight at −20 °C. The mixture precipitated overnight was centrifuged at 4°C, 12000×g for 30 min, the supernatant was discarded, and the ethanol was volatilized and dried. The precipitated protein was dissolved in lysate buffer (8 mol/L urea, 2 mol/L thiourea, 25 mmol/L dithiothreitol, 50 mmol/L Tris) and centrifuged at 4°C at 12000×g for 30 min, and the supernatant was placed in a new EP tube. The protein concentration was measured by Bradford method.

Enzyme digestion of urine protein: 100 μg urine protein sample was taken into 1.5 mL EP tube, and 25 mmol/L NH_4_HCO_3_ solution was added to make the total volume 200 μL. 20 mM Dithiothreitol solution (DTT, Sigma, prepared in 25 mmol/L NH4HCO3 solution) was added and mixed. The metal bath was heated at 97°C for 10 min, and cooled to room temperature. 50 mM Iodoacetamide (IAA, Sigma, prepared in 25 mmol/L NH4HCO3 solution) was added, mixed and reacted at room temperature for 40 min away from light. 200 μL UA solution (8 mol/L urea, 0.1 mol/L Tris-HCl, pH 8.5) was added to the 10 kDa ultrafiltration tube (Pall, Port Washington, NY, USA), Centrifuged at 18°C, 14000×g for 5 min and repeated once; freshly treated samples were added and centrifuged at 18□, 14000×g for 30 min, The urine protein was on the filter membrane. 200 μL UA solution was added, centrifuged at 18°C, 14000×g for 30 min and repeated twice. 25 mmol/L NH_4_HCO_3_ solution was added, centrifuged at 18°C, 14000×g for 30 min and repeated twice. Trypsin Gold (Promega, Fitchburg, WI, USA) was added at the ratio of 1:50 for enzyme digestion, and the water bath at 37°C for 15 h. After digestion, the peptides were collected by centrifugation at 4□ at 13000×g for 30 min, desalted by HLB solid phase extraction column (Waters, Milford, MA). The peptides were lyophilized with a vacuum dryer, and stored at -20°C.

#### 2.2.4 LC-MS/MS analysis

The digested samples were reconstituted with 0.1% formic acid, and peptides were quantified with BCA peptide quantification kit. The peptide concentration was then diluted with 0.1% formic acid to 0.5μg/μL. Mixed peptide samples were prepared from 6 μL of each sample and separated by high pH reversed phase peptide separation kit (Thermo Fisher Scientific, Waltham, MA, USA). Ten fractions were collected by centrifugation, lyophilized with a vacuum dryer and reconstituted with 0.1% formic acid. iRT reagent (Biognosys, Switzerland) was added to each fraction and each digested sample with a volume ratio of sample: iRT of 10:1 to calibrate the retention times of the extracted peptide peaks.

For analysis, 1 μg of peptide from each fraction and each digested sample was loaded onto a trap column and separated on a reverse-phase C18 column (50 μm ×150 mm, 2 μm) using the EASY-nLC1200 HPLC system (Thermo Fisher Scientific, Waltham, MA, USA). Peptides were analysed with an Orbitrap Fusion Lumos Tribrid Mass Spectrometer (Thermo Fisher Scientific, Waltham, MA, USA).

To generate the spectrum library, Ten fractions were subjected to mass spectrometry in data-dependent acquisition (DDA) mode, and 10 raw files were generated. Mass spectrometry data were collected in high sensitivity mode. A complete mass spectrometric scan was obtained in the 350–1200 m/z range with a resolution set at 60,000. 10 raw files were imported into Proteome Discoverer software for library construction using Swiss-iRT and Uniprot-Rat databases (version 2.0, Thermo Scientific). 39 variable window Data Independent Acquisition(DIA) methods for DIA mode of each digested sample were set up according to the results of database construction.

Each digested sample was analysed using Data Independent Acquisition (DIA) mode. DIA mode was performed using the DIA method with 39 windows. After every 7 samples, a single DIA analysis of the pooled peptides was performed as a quality control.

#### 2.2.5 Quantitative analysis of Label-free DIA

The raw file of each digested sample collected in DIA mode was imported into Spectronaut Pulsar(Biognosys AG, Switzerland) for analysis. The abundances of peptide were calculated by summing the peak areas of the respective fragment ions in MS_2_. Protein intensities were summed from their respective peptide abundances to calculate protein abundances.

#### 2.2.6 Data analysis

The LC-MS/MS analysis technique was repeated 3 times for each sample, and the PG.Quantity average value for each identified protein was taken for statistical analysis. Five rats were used as biological replicates. Five urine samples on the mating day group were compared with five urine samples on the next after mating day group. The identified proteins were compared to screen for differential proteins. The differential protein screening conditions were as follows: PG.Quantity fold change (FC) ≥ 1.5 or ≤ 0.67 between groups and *P* value < 0.05 by two-tailed unpaired t test analysis. The differential proteins were analyzed by Uniprot website (https://www.uniprot.org/) and the relevant literature was searched in Pubmed database (https://pubmed.ncbi.nlm.nih.gov) to analyze the function of the differential proteins.

## 3 Experimental results and analysis

### 3.1 Comparison of urine proteome between the day after mating and the day after mating

#### 3.2.1 Differential protein

All of the samples were identified 925 proteins. The urine protein of the next day of mating was compared with that of mating day. The screening conditions for differential protein were FC≥1.5 or ≤0.67, and the two-tailed unpaired T-test was P<0.05. The results showed that 43 differential proteins could be identified on the day after mating compared with the day of mating. First, the identified differential proteins were analyzed by using STRING database for protein interaction, and the results were shown in Figure 1. Then the differential proteins were arranged in the order of FC from largest to smallest and retrieved by Uniprot. The results were shown in Table 1.

**Table 1.**
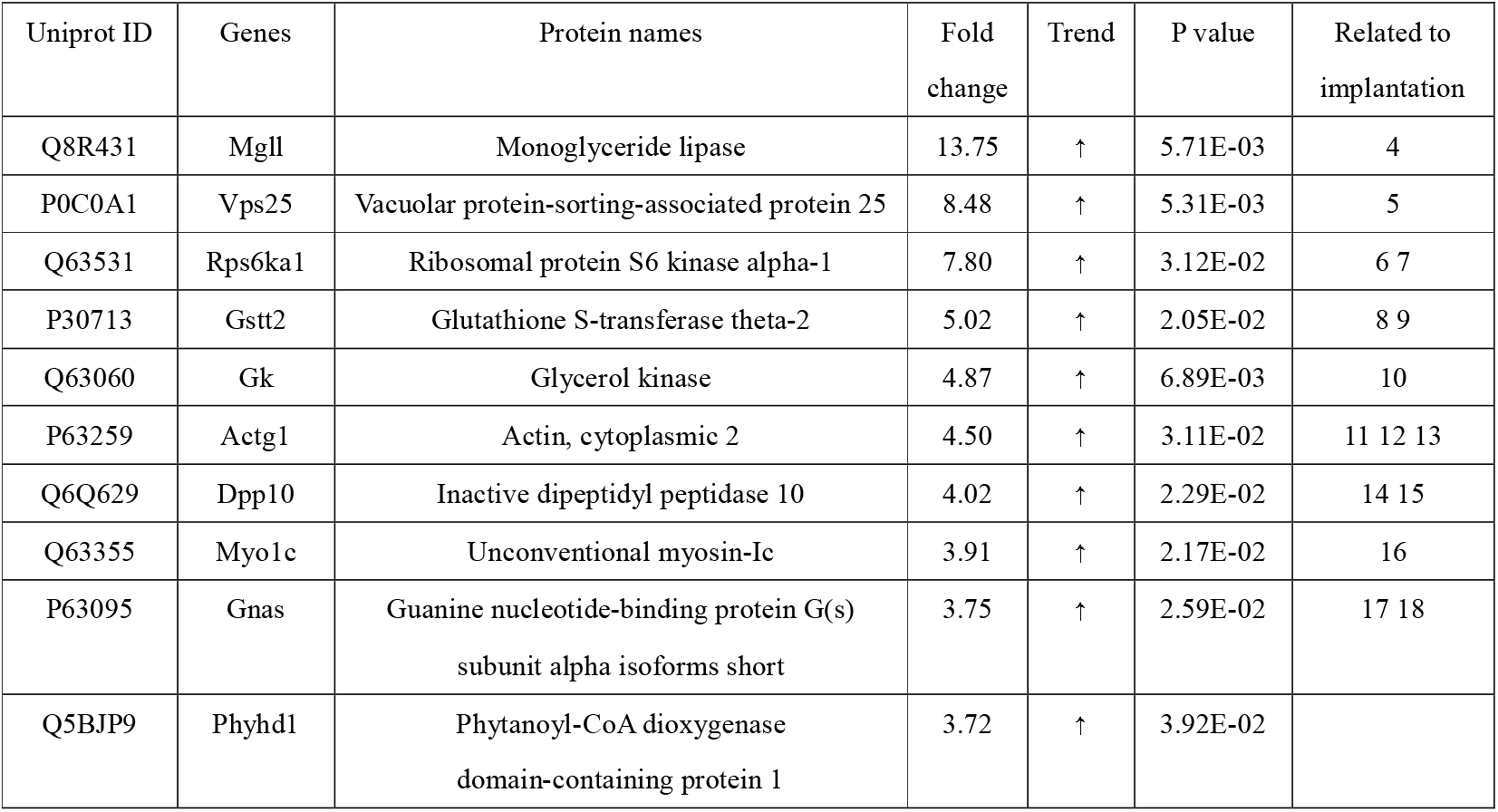
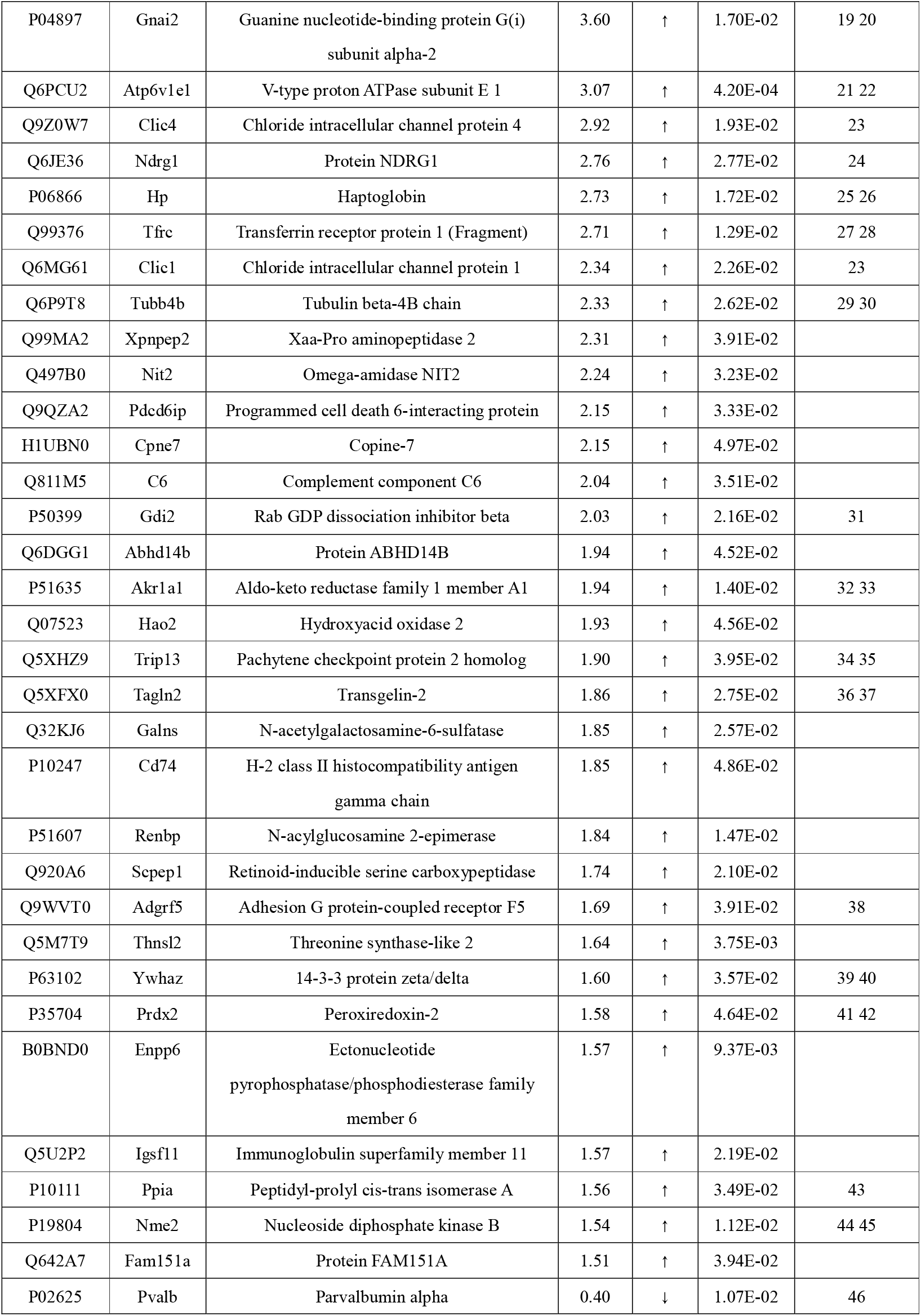
Urine differential proteins between the next day of mating and the day after mating.

**Figure 1.**
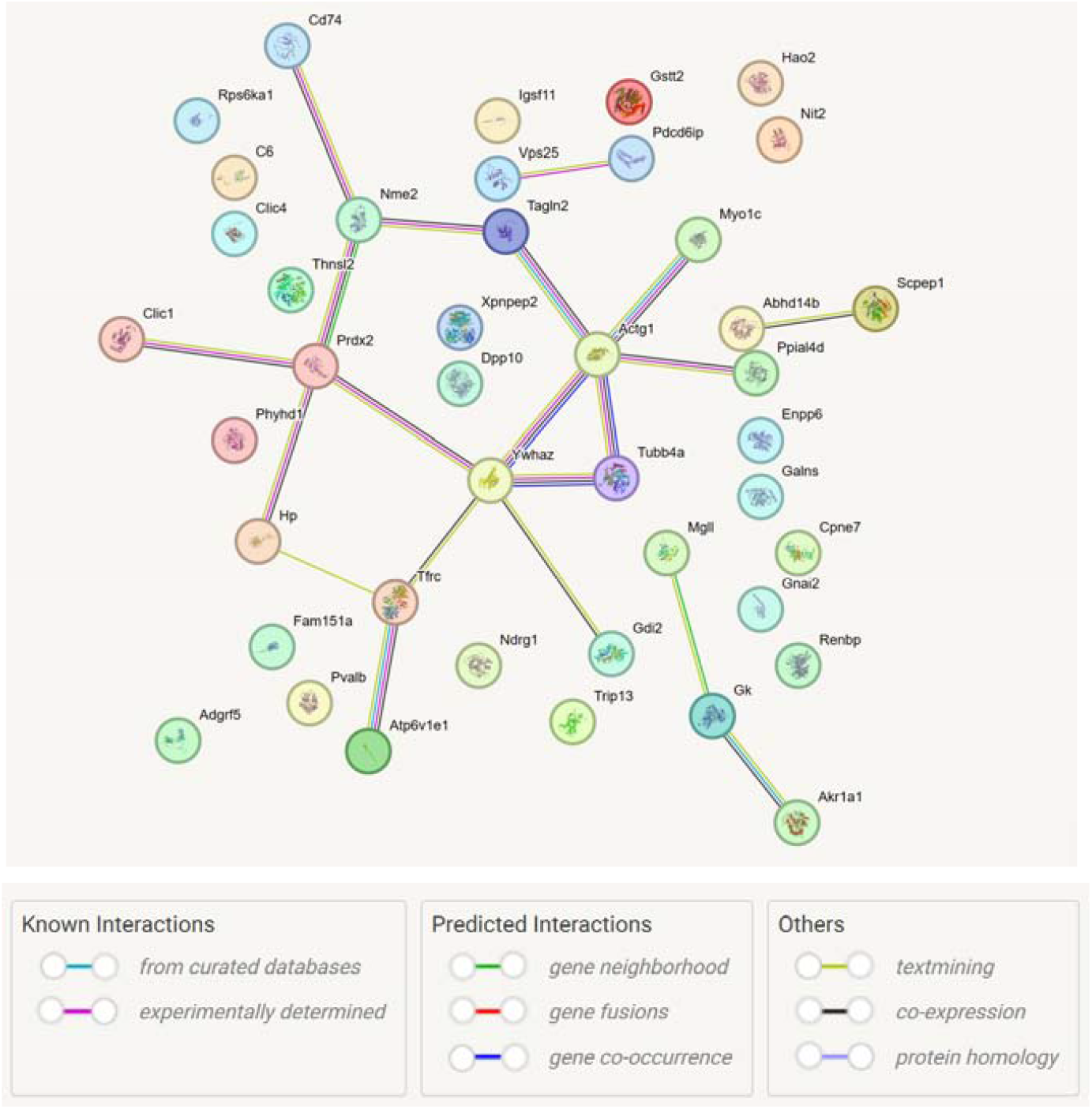
Interaction of urine differential proteins between the next day of mating and the day after mating

#### 3.2.2 Differential protein function analysis

The 43 identified differential proteins were searched through PubMed database, and 26 of them or other members of their family were reported to be associated with spermatogenesis. The function of these proteins is shown below:

Monoglyceride lipase is highly expressed in the testes and is directly involved in regulating human testicular physiology as part of the endocannabinoid system, including spermatogenesis and the function of stromal cells ^4^.

Mutations in Vacuolar protein-sorting associated protein 33 B from the same family as Vacuolar protein-sorting associated protein 25 cause sterility in C. elegans, Spermatocyte with division arrest indicates that this protein is involved in the formation of sperm-specific organelles ^5^.

Ribosomal protein S6 kinase alpha-1 substrate Ribosomal protein S6 is critical for spermatogenesis, and knockdown of this protein in Chinese bream resulted in spermatogenesis defects, including germ cell loss, mature sperm retention, and cavity formation ^6^. The protein can also regulate the blood-testicular barrier of supporting cells through Akt1/2, and then regulate the organization of F-actin, regulate the adhesion function of the intercellular interface, and promote the transport of blood-testicular barrier of spermatocytes in the pre-leptotene stage during rat spermatogenesis ^7^.

Glutathione S-transferase plays an important role in spermatogenesis and normal sperm function ^8^, The ineffective genotype of Glutathione S-transferase theta-1, the same family as Glutathione S-transferase theta-2, has been associated with dysspermatogenesis and may contribute to susceptibility to dysspermatogenesis and male infertility in the Chinese population ^9^.

Glycerol kinase 2, which has a high degree of homology with Glycerol kinase kinase, is crucial for the correct arrangement of crescent mitochondria to form mitochondrial sheath during mouse spermatogenesis. Knockout of this gene will lead to the disorder of mitochondrial sheath of sperm flagella ^10^.

Actin is involved in various aspects of spermatogenesis, and with spermatogenesis, the Actin cytoskeleton exhibits active remodeling during this process, participating in the shaping and differentiation of sperm cells ^11 12^. However, the molecular mechanisms by which Actin cytoskeletal tissue responds to spermatogenesis in spermatogenic epithelial cells remain largely unexplored ^13^.

The prostate secretes enzyme-containing vesicles called prost glands into human semen. At neutral or slightly acidic pH, the prost gland fuses with the sperm, thereby transferring certain molecules into the sperm. Dipeptidyl peptidase □ activity, which was not present in sperm, was transferred from the prost gland to sperm, allowing sperm to acquire a new membrane binding enzyme and change the catalytic activity of its surface ^14^. Guinea pig sperm contains Dipeptidyl peptidase II, and the protein is confined to a single compartment in the acrosomal sperm ^15^.

During spermatogenesis, spermatogonial cells undergo mitosis and meiosis to form bundles of sperm cells that remain connected to each other via cytoplasmic Bridges, which are individualized to form individual sperm cells. The process of individuation involves the formation of cytoskeletal proteins and membrane complexes around the sperm nucleus, the individualised complex, which breaks down the shared membrane into a single membrane surrounding each sperm cell. Unconventional myosin 95F myosin, which belongs to the Unconventional myosin family as well as Unconventional Myosin-IC, is a component of the individualization complex and is involved in membrane recombination during individualization. Its function is critical to the individualization process, and mutations in partial loss of function of this protein can lead to male infertility^16^.

Guanine nucleotide-binding protein G(s) subunit alpha is expressed in a tissue-specific and age-dependent manner in the reproductive organs of the ram. The protein is expressed at high levels in the epididymis, suggesting that it may influence the composition of the epididymal lumen fluid, thus, it affects the microenvironment of sperm maturation and may play an important role in spermatogenesis and the development of the testis and epididymis in the ram reproductive system ^17^. Guanine nucleotide-binding protein G(o) subunit alpha, from the same family as Guanine nucleotide-binding protein G(s) subunit alpha, The high level of expression in rat spermatocytes during pachytene stage suggests that it may play a role in this stage of spermatogenesis ^18^.

Guanine nucleotide-binding protein G(i) subunit alpha-1, Guanine nucleotide-binding protein G(i) subunit alpha-2, Guanine nucleotide-binding protein G(i) subunit alpha-3 and Guanine nucleotide-binding protein G(o) subunit alpha were detected in mouse spermatocytes and sperm cells. When spermatocytes develop into sperm cells, Guanine nucleotide-binding protein G(o) subunit alpha levels are reduced ^19^. Guanine nucleotide-binding protein G(i) subunit is associated with the developing acrosome, which may play a role in acrosome biogenesis. The Guanine nucleotidebinding protein G(i) subunit is found in the acrosomal region of mammalian sperm and is part of a complex required for signal transduction to induce acrosomal exocytosis ^19 20^.

V-type proton ATPases play an important role in the capacitation process of rabbit sperm ^21^. Rat round sperm cells regulate intracellular pH through V-type proton ATPases, HCO3- entry pathways, Na+/HCO3- dependent transport systems, and postulated proton conduction pathways. These pH adjustments appear to be specifically designed to withstand the acid challenge ^22^.

There are Chloride intracellular channel protein in bovine epididymis sperm_ο_ Chloride intracellular channel protein 1, Chloride intracellular channel protein 4 and Chloride intracellular channel protein 5, are present in the sperm, and occupy different positions in the cell. Both are able to bind to PP1γ2 in sperm, and since PP1γ2 is a key enzyme regulating sperm motility, the PP1γ2-binding protein Chloride intracellular channel protein may play an important role in sperm function^23^.

The N-myc downstream regulatory gene (NDRG) family consists of NDRG-1, NDRG-2, NDRG-3 and NDRG-4 members. It has been reported that Ndrg3 is a key gene that causes homologous lethal in early embryonic development, regulating male meiosis in mice. NDRG3 expression was specifically enhanced in germ cells and reached a peak level in spermatocytes of pachytene stage.In NdrG3-deficient germ cells, ERK activation is attenuated and double-strand break repair and synapsis complex formation in meiosis are impaired ^24^.

Haptoglobin, an iron transporter expressed in Sertoli, Leyden, and germ cells of the rat testis but not in the epididymis, may play an important role in iron metabolism in the testis. Testis Haptoglobin mRNA levels steadily increase during maturation after birth, indicating its involvement in spermatogenesis ^25^. Sertoli cells play a key role in spermatogenesis and express receptors for follicle-stimulating hormone (FSH) and testosterone (T), the major hormonal regulators of spermatogenesis.FSH stimulation of Sertoli cells in pigs resulted in an increase in inhibin-α, inhibin-β, plaquoglobin, haptoglobin, D-3-phosphoglycerate dehydrogenase, and sodium/potassium transport atpase in Sertoli extracellular vesicles ^26^.

The Transferrin receptor protein was identified in the proteome of newborn mouse testis, suggesting that it is involved in meiosis. Transferrin receptor proteins are essential for the progression of spermatocyte meiosis, particularly for DNA double-strand break repair and synapsis ^27^. The Transferrin receptor protein is only found in human spermatocytes and early spermatocytes.In patients with sperm genetic disorders, Transferrin is always present in Sertoli cells, and Transferrin receptor protein is only found when spermatocytes are present. In human seminiferous tubules, Sertoli cells are dedicated to Transferrin production, storage, and storage. Spermatocytes and early sperm cells use Transferrin^28^.

Many processes in spermatogenesis depend on the dynamic changes of the cytoskeleton, the movement of organelles, and especially the regulation of microtubules. Data from transgenic mouse models suggest that the coordination of microtubule dynamics is critical for male fertility ^29^_ο_

During spermatogenesis, a structure called “nuage” appears and disappears as spermatogenic cells differentiate. nuage can be divided into four types: Irregularly Shaped Perinuclear Granule (ISPG), Intermitintermital Cement (IMC), and Satellite Body (SB), Chromatoid Body (CB). ISPG, IMC and SB were observed in pachytene spermatocytes, while CB was observed in round sperms. In rat round sperm cells, Tubulin beta is translated from mRNA stored in CB, and assembled with Tubulin alpha outside CB to form a structural unit of microtubules: αβ-heterodimer, to construct microtubules in the sperm flella^30^.

Rab GDP dissociation inhibitor beta and family member Rab GDP dissociation inhibitor alpha are involved in the organization of actin cytoskeleton and also regulate cell morphology and cell motility. The expression of this protein is decreased in patients with asthenospermia ^31^.

Increased ability to metabolize specific steroids during testicular development, such as increased expression of Aldo-keto reductase family 1 member C3, the same family member as Aldo-keto reductase family 1 member A1, during testicular development in domestic cats ^32^. The level of Aldo-Keto Reductase mRNA in silkworm testis is higher than that in other tissues, which plays an important role in the spermatogenesis of silkworm ^33^.

Pachytene checkpoint protein 2 homolog was searched by Uniprot database. The results showed that this protein is expressed in the male reproductive nucleus, and is involved in biological processes such as sperm development, sperm formation, male meiosis, synaptic complex, meiosis recombination, double-strand break repair, and meiosis recombination checkpoint signaling pathway. This protein plays a key role in chromosome recombination and chromosome structure development during meiosis.In the early stage of meiosis recombination, it mediates the non-crossed pathway and also influences the crossed and non-crossed pathways to effectively complete homologous chromosome association, which is necessary for the effective sex chromosome synapsis. Mouse Pachytene checkpoint protein 2 is required for recombination and normal higher-order chromosome structure during meiosis. Pachytene checkpoint protein 2 plays a potential role in the non-crossed repair of double-strand breaks in meiosis. The spermatocytes of male mice with a complete mutation in this gene can fully associate with their chromosomes. However, in the process of chromosome recombination, the loss of this leads to the failure of repair of double-strand breaks, cell death at the pachytene stage, and the lack of post-meiotic cells in the testicular tissue ^34 35^. In males, Pachytene checkpoint protein 2 is required for sex chromosome syndesmosis and the formation of sex bodies (transcription-silencing subnuclear domains formed by the X and Y chromosomes) ^35^.

Acute heat stress impels translation, protein folding and protein degradation processes in chicken testis, leading to apoptosis and interfering with spermatogenesis. After acute heat stress, Transgelin is upregulated in the testis to resist heat-induced damage ^36^. The expression level of Transgelin gene was low in testicular tissue of aged animals ^37^.

Adhesion G protein-coupled receptor A3, which belongs to the same family as Adhesion G protein-coupled receptor F5, is a known marker of spermatogonial stem cells, and 55% of mice that have this gene deleted, even with normal spermatogenesis and epididymal sperm count, they are infertile from puberty ^38^.

The 14-3-3 protein plays a key regulatory role in both mitosis and meiosis. In mice, 14-3-3 protein epsilon is essential for normal sperm function and male fertility ^39^. Mature spermatids need to be released from their attached sertoli cells. Identified proteins involved in the adhesion between sertoli cells and mature spermatids include 14-3-3 protein zeta/delta, which was only found in the tubular segment lysate during sperm release. But what exactly is its role in spermatogenesis, and how does it interact or influence other signal transduction pathways in the testes remain unknown ^40^.

Peroxiredoxin-2 has some antioxidant properties and may be involved in maintaining oxidative balance in the spermatogenic environment of mice ^41^. Peroxiredoxin-2 also maintained the normal development of reproductive cells in newborn rats ^42^.

Peptidyl-prolyl cis-trans isomerase A is upregulated in the testes of mice treated with environmental estrogen, which reduces male sperm count and leads to male infertility. However, the molecular mechanism of its effect on male infertility remains unclear ^43^.

Nucleoside diphosphate kinase B is distributed in the manchette microtubule structure of sperm (a transient microtubule structure in elongated sperm that plays an important role in nuclear enrichment and sperm tail formation). Nucleoside diphosphate kinase A is distributed instantaneously in the round sperm nucleus and asymmetrically in the cytoplasm of the elongated sperm nucleus base. Nucleoside diphosphate kinase subtypes may have specific functions in phosphate transfer networks involved in human spermatogenesis and flagellar motility ^44^. Nucleoside diphosphate kinase plays a key role in spermatogenesis by increasing the level of glutathione peroxidase 5 in mouse cells to eliminate reactive oxygen species ^45^.

The combination of injection of testosterone undecanoate and oral levonorgestrel enhanced the expression of Parvalbumin alpha and inhibited spermatogenesis. Parvalbumin alpha protects testicular cells from apoptosis and promotes cell survival, and may be an early molecular target for hormone-induced spermatogenesis inhibition ^46^.

## 4 Discussion

Rats were self-controlled in the experiment, and the interval between two urine collection was only one day, so as far as possible, the interference of individual differences and their own growth and development could be excluded. Therefore, even with a small sample size, the results of this study could preliminarily show that there were significant differences in the urine proteome of rats on the second day after mating and on the mating day, and most of the differences were related to spermatogenesis. Although the correlation between the remaining differential proteins and spermatogenesis had not been found in the database, the results of this study suggested that these proteins may still be related to spermatogenesis and could be further studied as target proteins of spermatogenesis. This study demonstrated the potential of urine proteome in studying spermatogenesis, and provided a method of urine proteomics for exploring the pathogenesis of abnormal spermatogenesis in males, discovering targets and new diagnostic methods. And the results of this study indicated that the study of spermatogenesis by semen actually interfered with the spermatogenesis process at the same time as the collection of semen for the study, while the urine proteomic method in our study could be carried out without affecting spermatogenesis. Further experiments may consider expanding the sample size of experimental animals or collecting clinical samples for research. At the same time, it also reflected the sensitivity of urine proteome and opened up a new field of urine exploration.

